# ISWI remodeler facilitates cBAF genomic binding to drive cell fate transition

**DOI:** 10.64898/2026.05.08.723508

**Authors:** Young-Kwon Park, Ji-Eun Lee, Arthur I. Skoultchi, David J. Picketts, Weiqun Peng, Kai Ge

## Abstract

The ISWI chromatin remodeler regulates nucleosome spacing using one of two ATPase subunits Snf2h (Smarca5) and Snf2l (Smarca1). While *Snf2h* stable knockout (KO) is known to markedly reduce genomic binding of CTCF, an architectural protein organizing the 3D genome, ISWI’s role in regulating genomic binding and function of lineage-determining transcription factors (LDTFs) during cell fate transition remains largely unclear. Using conditional KO mice and derived cells, we show Snf2h and Snf2l are partially redundant and are required for embryonic development of muscle and adipose tissue as well as myogenesis and adipogenesis in culture. Stable KO of ISWI impairs LDTF-stimulated cell differentiation and disrupts *de novo* binding of the myogenic LDTF MyoD and the cBAF chromatin remodeler. Surprisingly, acute depletion of ISWI leaves *de novo* MyoD binding landscape largely intact while disrupting MyoD-dependent recruitment of cBAF and CTCF, with minimal effects on constitutive genomic binding of cBAF and CTCF. Together, our findings identify ISWI as an important mediator connecting LDTF binding to cBAF recruitment and chromatin organization during cell fate transition.

**Bullet points:**

- ISWI ATPases Snf2h and Snf2l are partially redundant and essential for muscle and adipose development
- ISWI is required for MyoD, C/EBPα, and PPARγ-driven cell fate transition
- Stable KO of ISWI disrupts genomic binding of MyoD, while acute depletion does not
- Acute ISWI deletion disrupts MyoD-dependent, but not constitutive, genomic binding of cBAF and CTCF

## Introduction

Cell fate transition and the maintenance of cell identity are controlled by lineage-determining transcription factors (LDTFs). During development, LDTFs coordinate transcriptional programs necessary for the differentiation of specific cell types from their progenitors. For example, MyoD functions as a key LDTF that drives myogenic program, while C/EBPα/β and PPARγ are essential LDTFs for adipogenesis ^1^. LDTF binding sites are often embedded within a restrictive chromatin environment. Therefore, chromatin remodeling is required to overcome nucleosome barriers and facilitate the recruitment of TFs and transcription coactivators, thereby establishing cell type-specific genomic landscape.

Chromatin remodeling is mediated by ATP-dependent remodelers that modulate chromatin accessibility by displacing, mobilizing, or restructuring nucleosomes ^2^. Among these remodelers, SWI/SNF family, especially the canonical BAF (BRG1/BRM-associated factor; cBAF) ^3^, and ISWI (Imitation SWItch) family represent two major classes with distinct biochemical properties and regulatory functions ^4^.

The cBAF chromatin remodeler mainly catalyzes nucleosome ejection to generate accessible chromatin for TFs and transcription coactivators ^2^. cBAF contains one of two mutually exclusive ATPases Brg1 or Brm as the catalytic subunit, along with additional components such as SS18 and the cBAF-specific Arid1a ^3,5,6^. cBAF plays a critical role in cell differentiation and animal development. Whole body knockout (KO) of *Brg1*, *Ss18*, or *Arid1a* leads to embryonic lethality in mice ^7–9^. Genetic deletion, knockdown or dominant-negative inhibition of cBAF impairs myogenesis and adipogenesis ^3,10,11^. cBAF facilitates cell differentiation by promoting enhancer activation and cell type-specific gene induction ^3,12–14^.

ISWI remodeler promotes nucleosome sliding and spacing to establish regularly ordered chromatin architecture ^4,15–18^. Mammalian ISWI contains either one of the two paralogous nucleosome-dependent ATPases Snf2h (Smarca5) and Snf2l (Smarca1) as the catalytic subunit. Snf2h and Snf2l associate with various accessory subunits to regulate diverse functions, including DNA replication, DNA repair, and transcription ^19^. Although Snf2h and Snf2l are 86% identical at the protein sequence level and have the same domain structure, they exhibit distinct expression profiles and developmental requirements *in vivo* ^20^. *Snf2h* KO embryos die at the peri-implantation stage before embryonic day 7.5, while *Snf2l* KO mice are viable and fertile ^21,22^. Despite these insights, the role of ISWI in chromatin reorganization during later lineage-specific differentiation programs remains poorly understood.

Here, we address this gap by investigating the role of ISWI during cell fate transition using myogenesis and adipogenesis as model systems. Using conditional KO mice and derived progenitor cells, we show essential and partially redundant roles of Snf2h and Snf2l in embryonic development of muscle and brown adipose tissue (BAT), and demonstrate a cell-autonomous requirement for ISWI in myogenesis and adipogenesis. Rescue experiments with an enzyme-dead mutant further establish that chromatin remodeling activity of ISWI is indispensable for cell differentiation. By comparing stable KO with dTAG-mediated acute depletion, we mechanistically distinguish immediate requirements of ISWI from long-term consequences of its loss. While stable KO of ISWI leads to widespread reduction of LDTF, cBAF, and CTCF genomic binding, acute depletion reveals that ISWI is largely dispensable for initial LDTF binding but essential for LDTF-dependent genomic binding of cBAF and CTCF.

## Results

### ISWI ATPases Snf2h and Snf2l are partially redundant and required for embryonic development of muscle and brown adipose tissue in mice

To explore the role of ISWI chromatin remodeler in cell fate transition, we focused on the ATPase subunits, Snf2h and Snf2l, in myogenesis and adipogenesis. Snf2h and Snf2l are 86% identical at the protein sequence level, share the same domain structure, and contain a highly conserved ATPase core (Figure 1A). We generated Myf5-Cre-mediated *Snf2h* conditional KO (*Snf2h*^f/f^;*Myf5-Cre*) mice by crossing *Snf2h*^f/f^ ^23^ with *Myf5-Cre* mice. Myf5-Cre is selectively expressed in precursor cells that give rise to skeletal muscle and BAT ^24^. *Snf2h*^f/f^;*Myf5-Cre* mice were obtained at the expected Mendelian ratio at embryonic day 18.5 (E18.5) but died shortly after birth (P0.5), likely due to defects in thoracic muscles (Figure 1B-C). Histochemistry analysis of sagittal sections along the midline revealed a significant reduction in muscle mass but a moderate decrease in BAT mass in E18.5 *Snf2h*^f/f^;*Myf5-Cre* embryos compared with littermate controls (Figure 1D). Deletion of *Snf2h* increased *Snf2l* expression by 2-fold in BAT, suggesting that Snf2l may partially compensate for the loss of *Snf2h in vivo* (Figure 1E).

**Figure 1.**
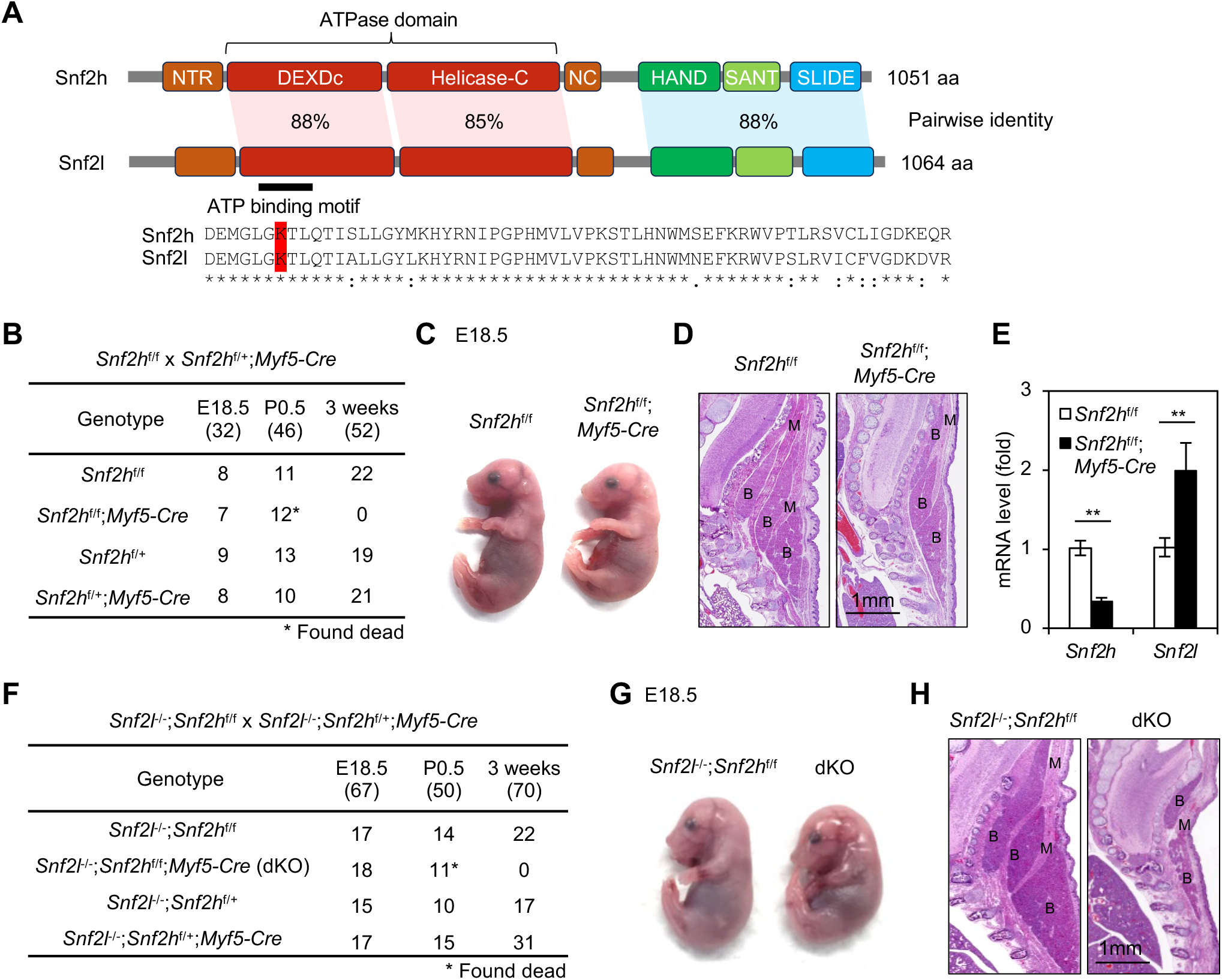
Snf2h and Snf2l are partially redundant and required for embryonic development of muscle and brown adipose tissue in mice. (**A**) Diagram of mouse Snf2h and Snf2l proteins. The locations of the ATPase domains and HAND–SANT–SLIDE domains, along with their degree of homology, are indicated. The lysine residue (K), essential for catalytic activity in the ATP-binding motif, is highlighted in red. (**B-E**) Snf2h is required for the embryonic development of muscle and brown adipose tissue (BAT) in mice. (**B**) Genotypes of progeny from crossing between *Snf2h*^f/f^ and *Snf2h*^f/+^;*Myf5-Cre* mice. (**C**) Representative images of E18.5 embryos of the indicated genotypes. (**D**) Sagittal sections of the cervical/thoracic area of E18.5 embryos. Embryos were stained with H&E. B, BAT; M, muscle. Scale bar = 1mm. (**E**) Increased expression of *Snf2l* in BAT of *Snf2h*^f/f^;*Myf5-Cre* embryos. RNA was extracted from E18.5 BAT of *Snf2h*^f/f^ (*n* = 4) or *Snf2h*^f/f^;*Myf5-Cre* (*n* = 4) embryos for qRT-PCR analysis. Statistical comparison between groups was performed using Student’s t-test (^∗∗^p < 0.05). (**F-H**) ISWI is required for adipose tissue and muscle development in mice. (**F**) Genotypes of progeny from crossing between *Snf2l*^-/-^;*Snf2h*^f/f^ and *Snf2l*^-/-^;*Snf2h*^f/+^;*Myf5-Cre* mice. (**G**) Representative images of E18.5 embryos of the indicated genotypes. (**H**) Sagittal sections of the cervical/thoracic area of E18.5 embryos.

To eliminate the compensatory effect, we crossed *Snf2l* KO (*Snf2l*^-/-^) and *Snf2h*^f/f^ mice and obtained *Snf2l/Snf2h* double conditional KO (*Snf2l*^-/-^;*Snf2h*^f/f^;*Myf5-Cre*, dKO) mice. *Snf2l*^-/-^ mice have been shown to survive without obvious phenotypes ^22^. dKO mice showed more severe defects in muscle and BAT development compared to *Snf2h* single KO at E18.5 and immediately died after birth (Figure 1F-H). These results indicate that ISWI ATPases Snf2h and Snf2l function in a partially redundant manner to promote development of muscle and BAT.

### ISWI ATPases Snf2h and Snf2l are essential for myogenesis and adipogenesis in culture

To validate the role of ISWI in myogenesis in culture, we isolated muscle satellite cells from 4OHT-inducible *Snf2h* conditional KO mice (*Snf2h*^f/f^;*CreER*) and induced myogenesis for 3 days. Deletion of *Snf2h* impaired myogenesis and prevented the induction of myocyte marker genes such as *Myog*, *Myh1*, and *Ckm* (Figure 2-S1A-B). Consistent with these results, shRNA-mediated stable knockdown of *Snf2h* led to severe defects in myogenesis of C2C12 myoblasts (Figure 2-S2A-C). Similarly, CRISPR-mediated KO of *Snf2h* in C2C12 cells did not alter MyoD expression (Figure 2-S2D-E) but resulted in defective myogenesis (Figure 2-S2F).

We also isolated muscle satellite cells from *Snf2l*^-/-^;*Snf2h*^f/f^;*CreER* mice and achieved simultaneous deletion of both *Snf2l* and *Snf2h* (hereafter referred to as ISWI KO). ISWI KO cells showed similarly severe defects in myogenesis (Figure 2A-B), indicating that myogenesis requires ISWI and is primarily mediated by Snf2h. To study ISWI-dependent transcription programs during myogenesis, we conducted RNA-Seq analyses in satellite cells before (day 0, D0) and after differentiation (day 3, D3). Using a 2-fold cutoff for differential expression, we identified 479 genes upregulated in a Snf2h-dependent manner, many of which were functionally associated with muscle cell differentiation (Figure 2C-D). During myogenesis, muscle satellite cells transition into myogenic precursor cells (myoblasts) before terminal differentiation into multinucleated myotubes (Figure 2E upper panel) ^25^. Notably, expression levels of *Pax7*, a TF essential for satellite cell maintenance, as well as early myogenic TFs *Myf5*, *Myod1*, *Myog*, and *Myf6*, were mostly comparable between ISWI KO and control cells. In contrast, genes involved in late-stage differentiation and myotube formation, including *Myh1*, *Myh2*, *Myh3*, *Erbb3*, and *Ncam1*, were significantly downregulated in ISWI KO cells (Figure 2E lower panel), suggesting that ISWI is largely dispensable for early myogenic commitment but required for terminal differentiation of muscle satellite cells.

**Figure 2.**
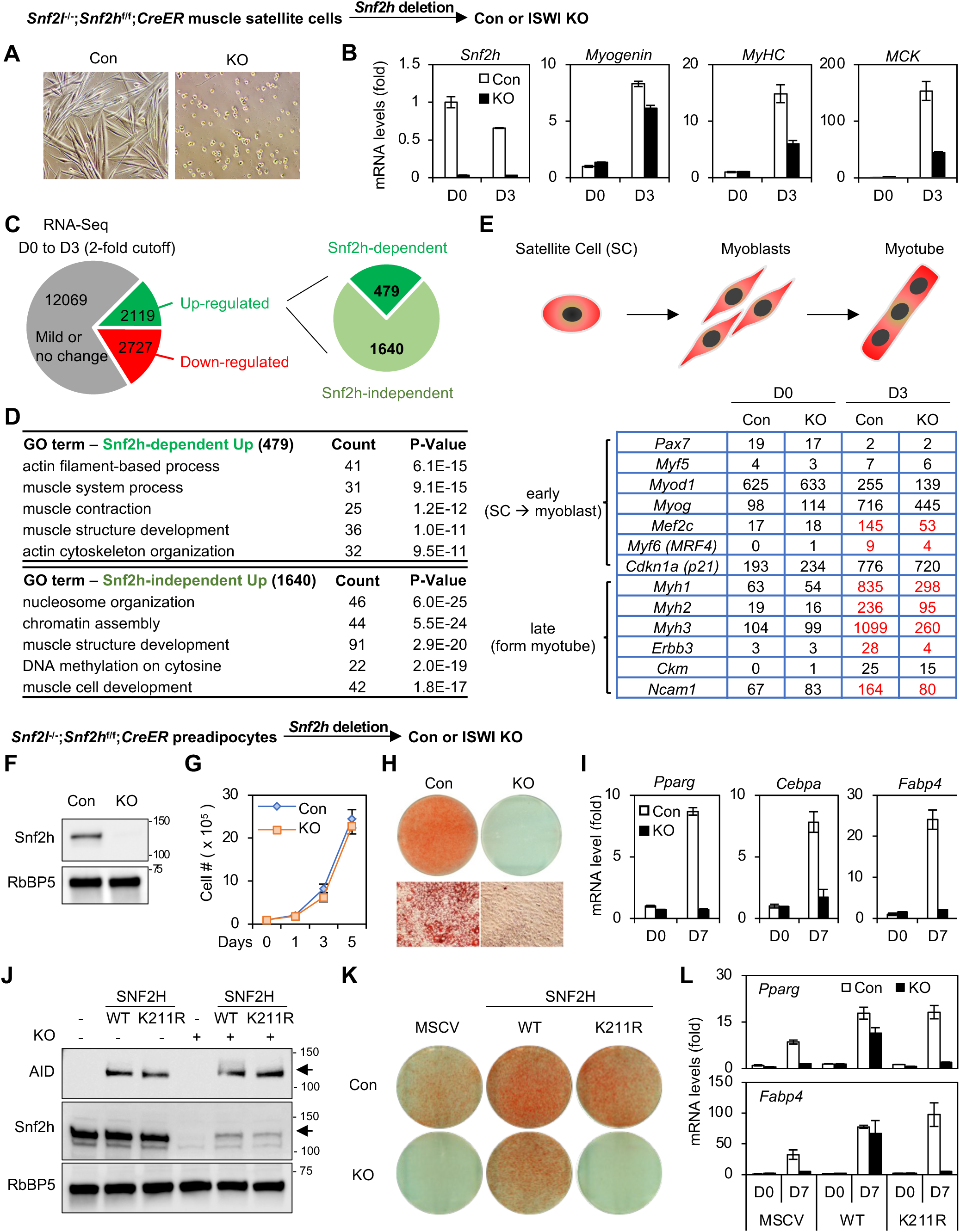
Snf2h and Snf2l are required for myogenesis and adipogenesis. (**A-E**) ISWI is required for myogenesis in culture. Muscle satellite cells isolated from *Snf2l*^-/-^;*Snf2h*^f/f^;*CreER* mice were treated with 4OHT to delete *Snf2h*, followed by myogenesis assays. (**A**) Representative microscopic pictures of satellite cells at D3 of myogenesis. (**B**) qRT-PCR analysis of myocyte marker genes *Myog*, *Myh1*, and *Ckm* during myogenesis. (**C**) RNA-Seq analysis before (D0) and after (D3) myogenesis in satellite cells. Pie chart depicts Snf2h-dependent and -independent up-regulated genes from D0 to D3 of myogenesis. The cut-off for differential expression is 2-fold. (**D**) GO analysis of 479 Snf2h-dependent and 1640 Snf2h-independent up-regulated genes defined in (**C**). (**E**) Expression of myogenic regulatory genes at D0 and D3 of myogenesis. Schematic diagram of satellite cell myogenesis (upper panel) and RPKM values (lower panel). >2-fold down-regulated genes in ISWI KO are highlighted in red. (**F-I**) ISWI is required for adipogenesis in culture. SV40T-immortalized *Snf2l*^-/-^;*Snf2h*^f/f^;*CreER* preadipocytes were treated with 4OHT, followed by adipogenesis assays. (**F**) Western blot analysis of Snf2h in preadipocytes. RbBP5 was used as a loading control. (**G**) Deletion of ISWI does not affect cell growth rates. 1 x 10^5^ preadipocytes were plated and cumulative cell numbers were determined for 5 days. (**H**) Oil Red O staining at D7 of adipogenesis. (**I**) qRT-PCR analysis of adipocyte marker genes *Pparg*, *Cebpa*, and *Fabp4* before (D0) and after (D7) adipogenesis. (**J-L**) ISWI enzymatic activity is required for adipogenesis. *Snf2l*^-/-^;*Snf2h*^f/f^;*CreER* cells were infected with retroviruses expressing AID-tagged wild type (WT) or enzyme dead mutant (K211R) of SNF2H, followed by 4OHT treatment to delete endogenous *Snf2h*. (**J**) Western blot analysis of Snf2h in preadipocytes. Arrows indicate ectopic SNF2H. (**K**) Oil Red O staining at D7 of adipogenesis. (**L**) qRT-PCR analysis of *Pparg* and *Fabp4* during adipogenesis.

Next, we investigated whether ISWI is also required for adipogenesis in culture. Preadipocytes isolated from BAT of *Snf2h*^f/f^;*CreER* and *Snf2l*^-/-^;*Snf2h*^f/f^;*CreER* mice were immortalized and treated with 4OHT to delete *Snf2h*. Deletion of *Snf2h* alone or simultaneous deletion of both *Snf2l* and *Snf2h* did not affect cell proliferation (Figure 2-S1C-D and Figure 2F-G), but completely blocked adipogenesis and the induction of adipocyte marker genes, including *Pparg*, *Cebpa*, and *Fabp4* (Figure 2-S1E and Figure 2H-I). Moreover, shRNA-mediated stable knockdown of *Snf2h* similarly led to severe defects in adipogenesis of preadipocytes (Figure 2-S2G-I). RNA-Seq analysis revealed that *Snf2h* was abundantly expressed before (day -3, D-3) and during (day 2, D2) adipogenesis, whereas *Snf2l* expression remained low and was not upregulated upon *Snf2h* deletion (Figure 2-S1F-G). Consistent with the morphological defects, RNA-Seq analysis confirmed that Snf2h-dependently upregulated genes were functionally associated with fat cell differentiation (Figure 2-S1H-I).

To determine whether ISWI chromatin remodeling activity is required for cell differentiation, we re-introduced wild-type (WT) or enzyme-dead mutant SNF2H (SNF2H^K211R^) ^26^ into ISWI KO preadipocytes. While WT SNF2H rescued adipogenesis, mutant SNF2H failed to do so, demonstrating that the ISWI chromatin remodeling activity is required for adipogenesis (Figure 2J-L). Together, these data indicate that ISWI is essential for myogenesis and adipogenesis in culture.

### Stable KO of ISWI prevents LDTF-stimulated myogenesis and adipogenesis

We next asked whether ISWI is required for cell fate transition driven by LDTFs. Ectopic expression of the myogenic LDTF MyoD is sufficient to convert preadipocytes into myocytes ^1^. To test whether ISWI is necessary for MyoD-driven cell fate transition, we generated preadipocytes expressing doxycycline (Dox)-inducible T7-tagged MyoD (Figure 3A). Induction of MyoD successfully converted control preadipocytes into myocytes and upregulated myocyte-enriched genes, including *Myog*, *Myh1*, and *Ckm*. In contrast, stable KO of ISWI impaired MyoD-stimulated myogenesis and failed to induce myocyte gene expression (Figure 3B-C).

**Figure 3.**
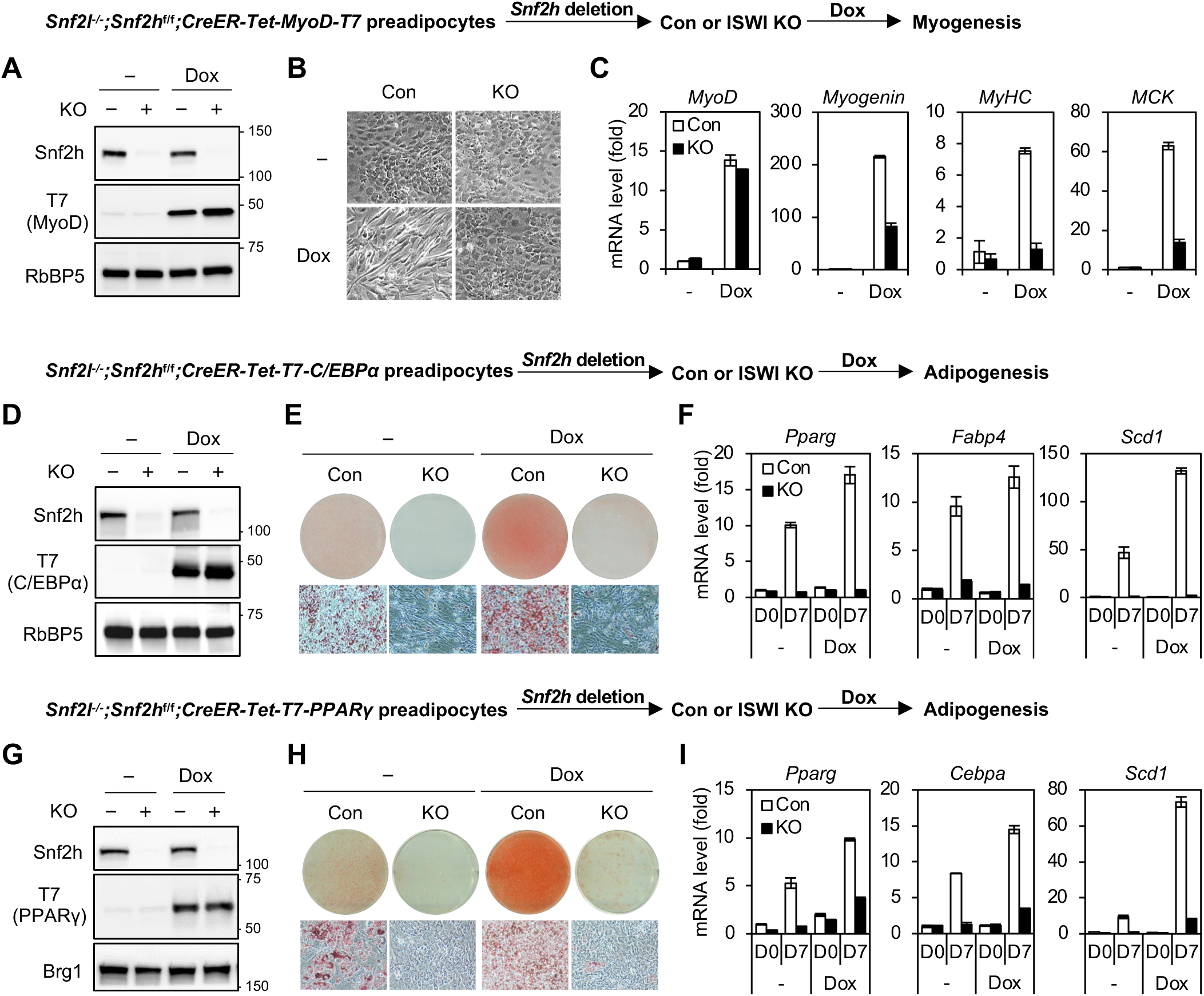
Stable KO of ISWI prevents LDTF-stimulated myogenesis and adipogenesis. (**A-C**) ISWI is required for MyoD-mediated myogenesis. *Snf2l*^-/-^;*Snf2h*^f/f^;*CreER* preadipocytes were infected with lentiviruses expressing doxycycline (Dox)-inducible MyoD-T7. Cells were treated with 4OHT to delete *Snf2h*, followed by myogenesis assay in the presence of Dox for 5 days. (**A**) Western blot analysis of Snf2h and Dox-induced MyoD-T7 expression. (**B**) Microscopic pictures at D5 of myogenesis. (**C**) qRT-PCR analysis of ectopic *MyoD* and myocyte-enriched genes *Myog*, *Myh1*, and *Ckm* at D5 of myogenesis. (**D-I**) ISWI is required for C/EBPα- and PPARγ-stimulated adipogenesis. *Snf2l*^-/-^;*Snf2h*^f/f^;*CreER* preadipocytes were infected with lentiviruses expressing Dox-inducible T7-C/EBPα or T7-PPARγ. Cells were treated with 4OHT to delete *Snf2h*, followed by adipogenesis assay in the presence of Dox for 7 days. (**D** and **G**) Western blot analysis of Snf2h and Dox-induced T7-C/EBPα or T7-PPARγ expression. RbBP5 was used as a loading control. (**E** and **H**) Oil Red O staining at D7 of adipogenesis. (**F** and **I**) qRT-PCR analysis of adipocyte marker genes *Pparg*, *Cebpa*, *Fabp4*, and *Scd1* at D0 and D7 of adipogenesis.

We next examined whether ISWI is similarly required for adipogenic cell fate transition driven by the adipogenic LDTFs C/EBPα and PPARγ. Dox-induced C/EBPα levels were comparable between control and ISWI KO cells (Figure 3D). While ectopic C/EBPα enhanced lipid accumulation and adipocyte marker gene expression in control cells, it failed to rescue adipogenesis defects in ISWI KO preadipocytes (Figure 3E-F). We also generated preadipocytes expressing Dox-inducible T7-tagged PPARγ (Figure 3G). Ectopic PPARγ failed to rescue adipogenesis defects in ISWI KO cells, regardless of stimulation with either a conventional adipogenic cocktail or the PPARγ agonist rosiglitazone (Rosi) (Figure 3H and Figure 3-S1A). Consistent with the morphological defects, ectopic PPARγ failed to induce adipogenic marker gene expression in ISWI KO cells (Figure 3I and Figure 3-S1B). Together, these findings demonstrate that ISWI is essential for LDTF-stimulated myogenic and adipogenic cell fate transition.

### Stable KO of ISWI disrupts *de novo* binding of MyoD and cBAF on chromatin

Having observed the failure of MyoD-stimulated myogenesis in ISWI KO cells, we examined whether ISWI is required for chromatin binding of MyoD and the chromatin remodeler cBAF, as cBAF is known to associate with MyoD and is required for myogenesis ^11^. Western blot analyses confirmed that expression levels of Dox-induced MyoD and the cBAF subunits were comparable in control and ISWI KO cells (Figure 4A).

**Figure 4.**
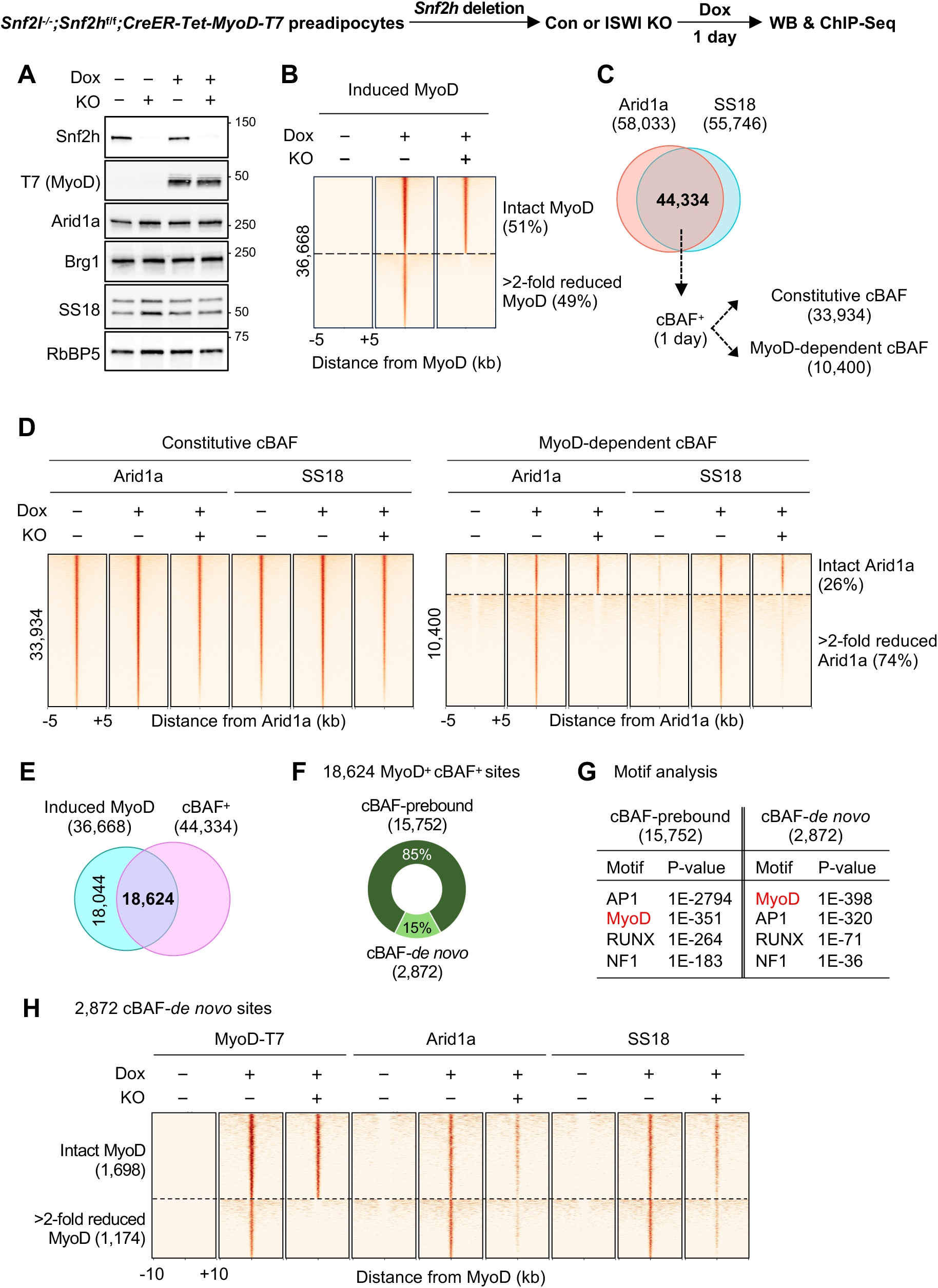
Stable KO of ISWI disrupts *de novo* binding of MyoD and cBAF on chromatin. *Snf2l*^-/-^;*Snf2h*^f/f^;*CreER* preadipocytes were infected with lentiviruses expressing Dox-inducible MyoD-T7 (MyoD). Cells were treated with 4OHT to delete *Snf2h*, followed by 1 day Dox treatment to induce MyoD expression. Sub-confluent cells were collected for ChIP-Seq analyses. (**A**) Western blot analysis of Snf2h, Dox-induced MyoD expression, and cBAF subunits Arid1a, Brg1, and SS18. RbBP5 was used as a loading control. (**B**) Stable KO of ISWI disrupts *de novo* binding of MyoD. Heat maps illustrate changes in MyoD binding intensity on 36,668 induced MyoD binding sites, grouped into intact MyoD (18,727) and >2-fold reduced MyoD (17,941) sites. (**C-D**) Stable KO of ISWI disrupts MyoD-dependent, but not constitutive, cBAF binding. (**C**) Venn diagrams depicting identification of cBAF^+^ sites by overlapping Arid1a and SS18 binding sites. cBAF^+^ sites were further grouped into constitutive or MyoD-dependent sites based on Arid1a binding status before inducing MyoD expression. (**D**) Heat maps illustrating changes in Arid1a and SS18 binding intensity on 33,934 constitutive cBAF binding sites or 10,400 MyoD-dependent cBAF binding sites. (**E-H**) Stable KO of ISWI disrupts *de novo* binding of MyoD and cBAF. (**E**) Venn diagrams depicting overlap between induced MyoD binding sites and cBAF^+^ regions. (**F**) Donut chart illustrating cBAF binding status on 18,624 MyoD^+^ cBAF^+^ sites identified in (**E**). **(G**) Motif analysis of cBAF-prebound and cBAF-*de novo* sites defined in (**F**). (**H**) Heat maps on 2,872 cBAF-*de novo* sites, grouped into intact MyoD (1,698) and reduced MyoD (1,174) sites. The cut-off for MyoD binding change is 2-fold.

We profiled global MyoD binding by ChIP-Seq using a T7 antibody. Among 36,668 *de novo* MyoD binding sites identified after 1 day of Dox-treatment, 49% showed a significant reduction (>2-fold) in MyoD signal intensity in ISWI KO cells (Figure 4B and Figure 4-S1A-C). We next asked whether ISWI loss also affects global cBAF chromatin binding. By overlapping Arid1a and SS18 ChIP-Seq peaks, we identified 44,334 high-confidence cBAF^+^ sites. These sites were further classified into constitutive or MyoD-dependent sites based on changes in Arid1a occupancy following MyoD induction. Constitutive cBAF sites (33,934) exhibit stable Arid1a binding before and after MyoD induction, whereas MyoD-dependent cBAF sites (10,400) show *de novo* binding of Arid1a upon MyoD induction (Figure 4C). Interestingly, while constitutive cBAF binding remained largely unchanged, MyoD-dependent *de novo* cBAF binding was significantly impaired in ISWI KO cells (Figure 4D).

To determine how changes in MyoD binding relate to cBAF chromatin occupancy, we focused on 18,624 sites co-occupied by MyoD and cBAF (Figure 4E). 85% of these MyoD^+^ cBAF^+^ sites exhibited cBAF binding before MyoD induction (referred to as cBAF-prebound), while the remaining 15% showed MyoD-dependent recruitment of cBAF (referred to as cBAF-*de novo*) (Figure 4F). Motif analysis revealed that cBAF-prebound sites show predominant enrichment of AP1 motifs. In contrast, the MyoD motif was the most significantly enriched one in cBAF-*de novo* sites, suggesting these sites likely represent primary MyoD targets during cell fate transition (Figure 4G). Consistent with the global analysis, MyoD and cBAF binding was significantly disrupted on these 2,872 cBAF-*de novo* sites in ISWI KO cells (Figure 4H and Figure 4-S1D). Interestingly, stable KO of ISWI impaired cBAF occupancy on both MyoD-intact and MyoD-reduced sites, suggesting that ISWI is fundamentally required for cBAF recruitment to MyoD^+^ sites regardless of the MyoD binding levels. Collectively, these results indicate that stable loss of ISWI results in pervasive disruption of *de novo* MyoD and cBAF binding during myogenic cell fate transition.

### Acute depletion of ISWI leaves *de novo* MyoD binding landscape largely intact

Because stable KO of ISWI may lead to secondary effects on chromatin landscape, we employed a dTAG-mediated acute depletion system. For this purpose, we introduced ectopic SNF2H-dTAG into *Snf2l*^-/-^;*Snf2h*^f/f^;*CreER* preadipocytes and deleted endogenous *Snf2h* to generate a cell line in which ISWI function is entirely dependent on the SNF2H-dTAG protein (Figure 5-S1). By combining this system with 4OHT-induced rapid MyoD-ER (hereafter referred to as MyoD) nuclear translocation, we could closely monitor the immediate consequences of ISWI depletion on *de novo* MyoD binding (Figure 5A). We confirmed that dTAG-13 treatment for 3h successfully depleted ISWI but did not affect the nuclear translocation of MyoD (Figure 5B). In contrast to the widespread defects observed in stable KO cells, acute depletion of ISWI resulted in only a modest decrease in *de novo* MyoD binding. Among 60,028 MyoD binding sites identified 1h after nuclear translocation, only 17% exhibited a significant reduction (>2-fold) in MyoD signal intensity upon ISWI depletion (Figure 5C). MyoD binding around the *Myog* and neighboring gene loci was not compromised by the acute depletion of ISWI (Figure 5D). These results suggest that the disrupted MyoD binding observed in stable KO cells was likely a secondary consequence of prolonged ISWI deficiency. To test this hypothesis, we examined the effects of 48h of ISWI depletion (Figure 5E-F). Indeed, prolonged ISWI depletion led to a progressive and significant reduction in MyoD binding (Figure 5G). Together, these findings suggest that ISWI is largely dispensable for *de novo* MyoD binding but is required for the long-term maintenance of the myogenic chromatin landscape.

**Figure 5.**
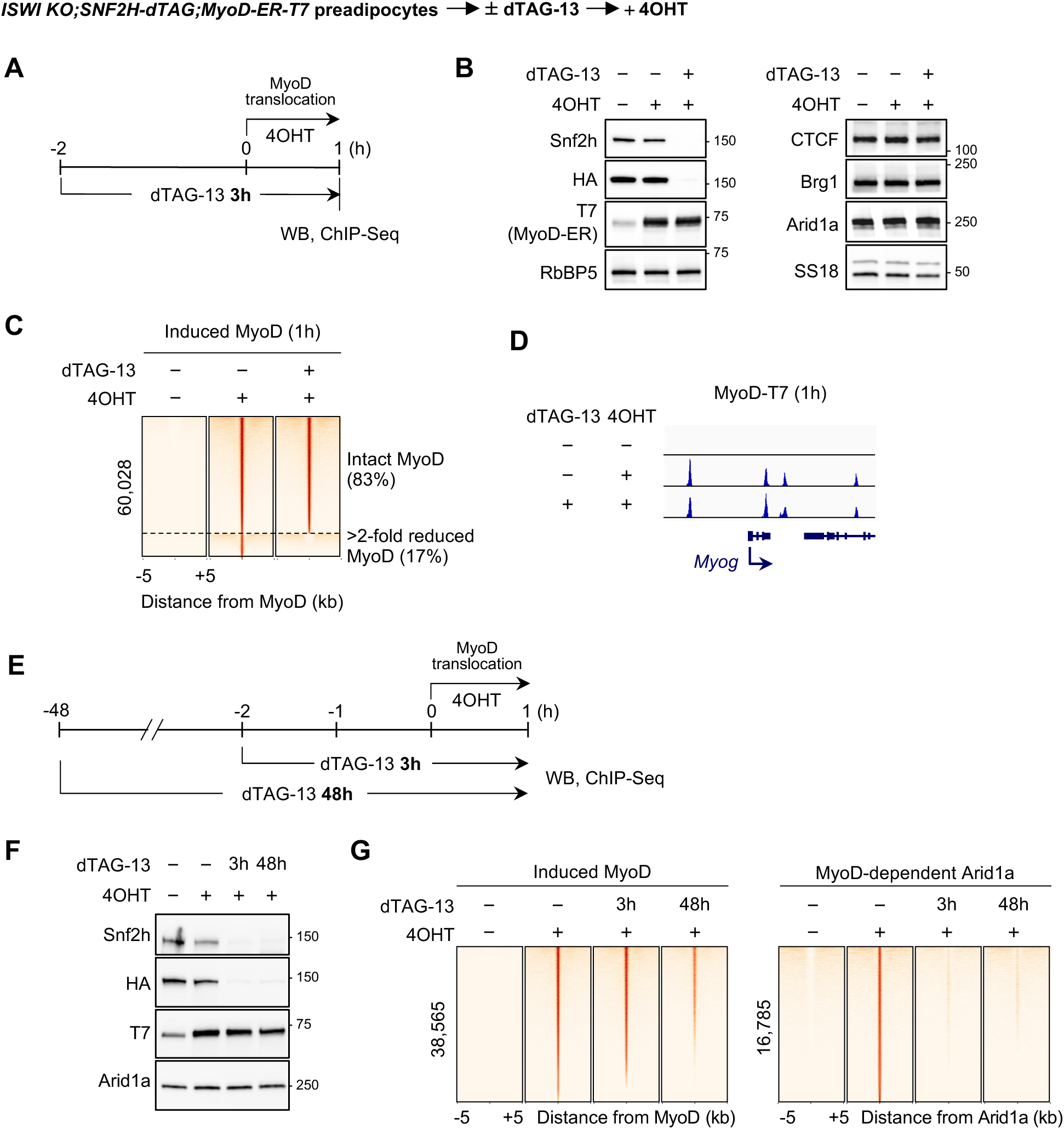
Acute depletion of ISWI leaves *de novo* MyoD binding landscape largely intact. *ISWI KO; SNF2H-dTAG* preadipocytes were infected with retroviruses expressing MyoD-ER-T7. Cells were pretreated with dTAG-13 to deplete SNF2H protein, followed by 4OHT treatment to induce MyoD nuclear translocation for an additional 1h. (**A-D**) Acute depletion of ISWI leaves *de novo* MyoD binding landscape largely intact. (**A**) Experimental schematic. (**B**) Western blot analysis of SNF2H-dTAG-HA depletion, MyoD nuclear translocation, CTCF, and cBAF subunits Arid1a, Brg1, and SS18 expression. RbBP5 was used as a loading control. (**C**) Heat maps illustrate changes in MyoD binding intensity on 60,028 induced MyoD binding sites, grouped into intact MyoD (49,649) and >2-fold reduced MyoD (10,379) sites. (**D**) Genome browser view of MyoD binding on *Myog* and neighboring gene loci. (**E-G**) Prolonged ISWI depletion leads to progressive reduction in *de novo* MyoD binding. (**E**) Experimental schematic. (**F**) Western blot analysis of SNF2H-dTAG-HA depletion and MyoD nuclear translocation. (**G**) Heat maps showing changes in MyoD and Arid1a binding intensity across ±5kb from the center of binding sites.

### Acute depletion of ISWI disrupts *de novo* cBAF binding to MyoD^+^ sites with little effects on constitutive cBAF binding

Since acute depletion of ISWI does not substantially affect *de novo* MyoD binding landscape, we next assessed its effects on cBAF chromatin binding. We first determined the global cBAF binding landscape by overlapping Arid1a and Brg1 ChIP-Sqe peaks 1h after inducing MyoD nuclear translocation. Among 29,711 cBAF^+^ sites, 48% (14,180) represented constitutive cBAF sites already bound by Arid1a before MyoD nuclear translocation. The remaining 52% (15,531) represented MyoD-dependent cBAF sites where Arid1a was recruited in response to MyoD nuclear translocation (Figure 6A). Consistent with our data in ISWI stable KO cells, acute depletion of ISWI had little effect on constitutive cBAF binding, but led to a significant reduction in MyoD-dependent cBAF binding (Figure 6B). Since nuclear translocated MyoD colocalizes with cBAF and the histone acetyltransferases CBP/p300 ^27^, we extended our analysis to CBP. Consistent with selective loss of MyoD-dependent cBAF binding, acute depletion of ISWI also significantly impaired MyoD-dependent CBP binding (Figure 6-S1A). These results support the role of ISWI in coordinating MyoD-induced chromatin reorganization.

**Figure 6.**
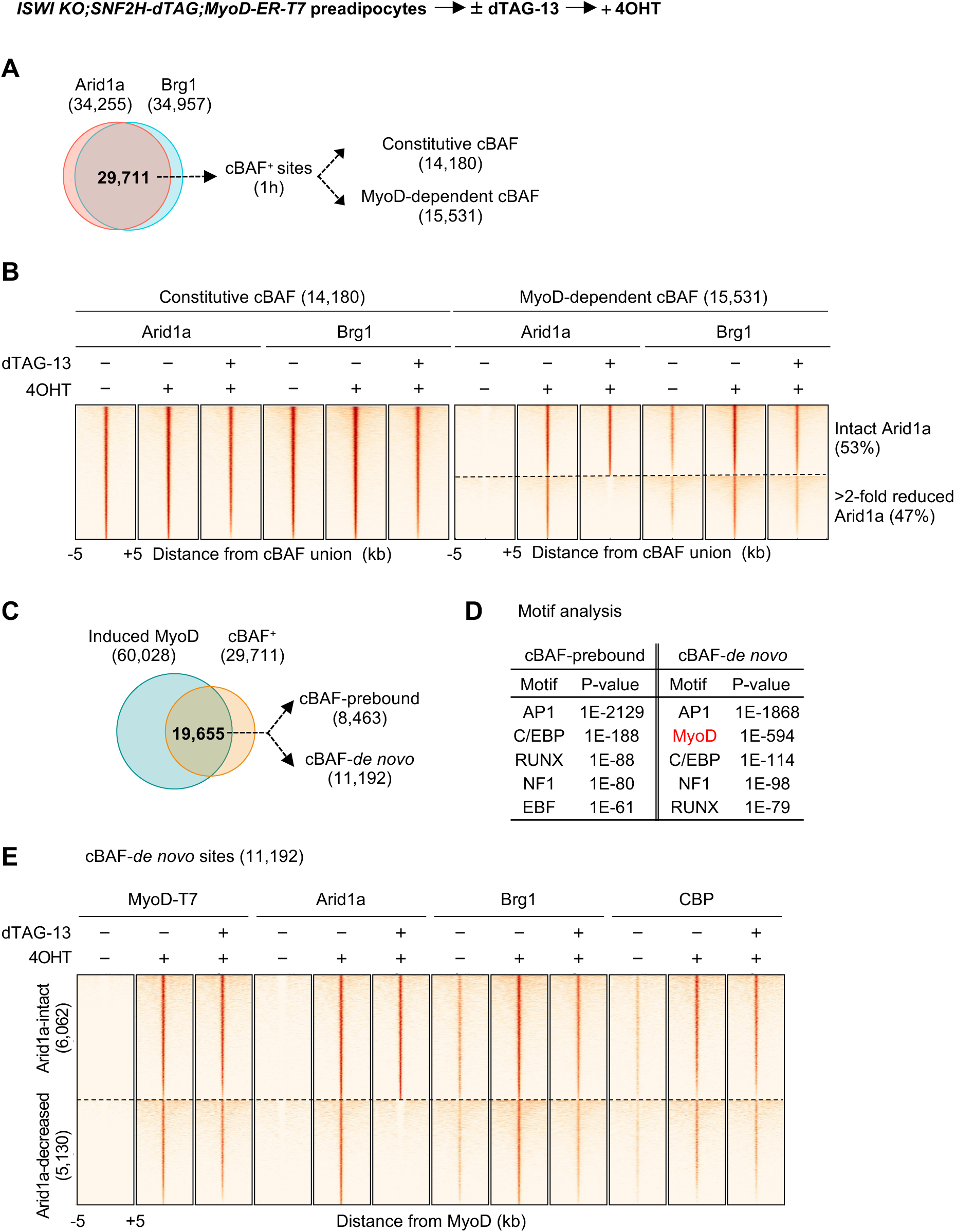
Acute depletion of ISWI disrupts *de novo* cBAF binding to MyoD^+^ sites with little effects on constitutive cBAF binding. *ISWI KO; SNF2H-dTAG* preadipocytes were infected with retroviruses expressing MyoD-ER-T7. Cells were pretreated with dTAG-13 for 2h to deplete SNF2H protein, followed by 4OHT treatment to induce MyoD nuclear translocation for an additional 1h. (**A-B**) Acute depletion of ISWI disrupts MyoD-dependent, but not constitutive, cBAF binding. (**A**) Venn diagrams depicting identification of cBAF^+^ sites by overlapping Arid1a and Brg1 binding sites. cBAF^+^ sites were further grouped into constitutive or MyoD-dependent sites based on Arid1a binding status before MyoD nuclear translocation. (**B**) Heat maps of 14,180 constitutive, and 15,531 MyoD-dependent, cBAF sites. MyoD-dependent cBAF sites grouped into intact Arid1a (8,287) and >2-fold reduced Arid1a (7,244) sites. (**C-E**) Acute depletion of ISWI disrupts *de novo* cBAF binding to MyoD^+^ sites. (**C**) Venn diagrams depicting overlap between induced MyoD and cBAF^+^ sites. The overlapped sites were further grouped into cBAF-prebound and cBAF-*de novo* sites based on Arid1a binding status before MyoD nuclear translocation. (**D**) Motif analysis of cBAF-prebound and cBAF-*de novo* sites defined in (**C**). (**E**) Heat maps illustrating changes in MyoD, Arid1a, Brg1 and CBP binding intensity on 11,192 cBAF-*de novo* sites.

We next focused on genomic regions occupied by both MyoD and cBAF. MyoD^+^ cBAF^+^ sites identified 1h after MyoD nuclear translocation separated into two distinct groups: cBAF-prebound sites (43%; 8,463), where Arid1a binding was already present before MyoD nuclear translocation, and cBAF-*de novo* sites (57%; 11,192), where Arid1a was newly recruited upon MyoD nuclear translocation (Figure 6C). Motif analysis revealed a significant enrichment of MyoD motifs exclusively within cBAF-*de novo* sites (Figure 6D), indicating that cBAF-*de novo* sites are primary targets of the MyoD-driven remodeling program. Therefore, we evaluated the effects of ISWI depletion on 11,192 cBAF-*de novo* sites. Consistent with global analysis, ISWI depletion significantly reduced both Arid1a and Brg1 binding without disrupting MyoD binding (Figure 6E and 6-S1B). The changes in CBP binding upon ISWI depletion were more heterogeneous, ranging from loss to gain.

Collectively, our data highlight a key distinction between ISWI stable KO and acute depletion. Acute depletion of ISWI primarily impairs *de novo* cBAF binding to MyoD^+^ sites during early cell fate transition, even when MyoD successfully engages its target sites.

### Acute depletion of ISWI disrupts MyoD-dependent but not constitutive CTCF binding on chromatin

Earlier studies using gene KO or knockdown approaches have identified ISWI as a critical regulator of CTCF genomic binding and nucleosome organization at CTCF sites ^26,28^. Consistent with these reports, we found that stable KO of ISWI in preadipocytes did not alter CTCF protein levels (Figure 7A), but resulted in a significant reduction in CTCF binding, with more than 50% showing >2-fold decrease in CTCF signal intensity (Figure 7B-C).

**Figure 7.**
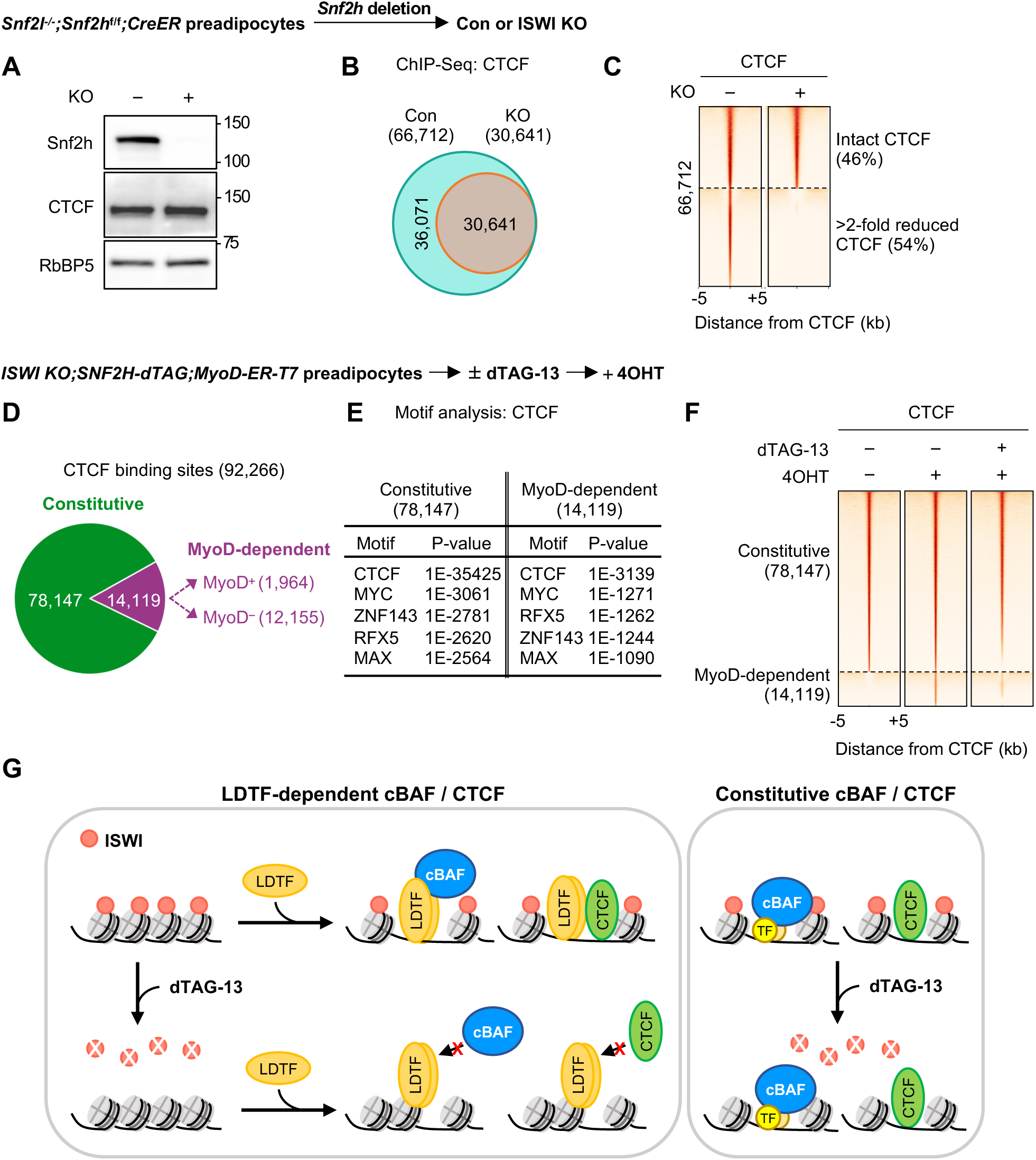
Acute depletion of ISWI does not affect constitutive CTCF binding but disrupts MyoD-dependent CTCF binding on chromatin. (**A-C**) Stable KO of ISWI reduces CTCF genomic binding. (**A**) Western blot analysis of Snf2h and CTCF. RbBP5 was used as a loading control. (**B**) Venn diagrams depicting CTCF binding sites in control and ISWI stable KO cells. (**C**) Heat maps illustrating changes in CTCF binding intensity. (**D-F**) Acute depletion of ISWI does not affect constitutive CTCF binding but disrupts MyoD-dependent CTCF binding. *ISWI KO; SNF2H-dTAG* preadipocytes were infected with retroviruses expressing MyoD-ER-T7. Cells were pretreated with dTAG-13 for 2h to deplete SNF2H protein, followed by 4OHT treatment to induce MyoD nuclear translocation for an additional 1h. Cells were then collected for ChIP-Seq analyses. (**D**) Pie chart illustrating classification of CTCF binding sites identified 1h after MyoD nuclear translocation. (**E**) Motif analysis of CTCF binding sites identified in (**D**). (**F**) Heat maps illustrating changes in CTCF binding intensity on 78,147 constitutive CTCF sites and 14,119 MyoD-dependent CTCF sites. (**G**) Working model. ISWI promotes LDTF-dependent genomic binding of cBAF and CTCF. Acute depletion of ISWI selectively disrupts LDTF-dependent, but not constitutive, genomic occupancy of the cBAF chromatin remodeler and the architectural factor CTCF.

Although the vast majority of CTCF binding sites are conserved across cell types ^29^, a subset is dynamically established during cell fate transition through cooperation with LDTFs. In myocytes, MyoD facilitates muscle-specific CTCF binding to control muscle cell identity ^30^. Given our observation that stable KO and acute depletion of ISWI exert distinct effects on MyoD genomic binding (Figures 4 and 5), we next asked whether loss of CTCF binding in stable KO cells reflects a primary consequence of ISWI depletion or a secondary effect of long-term chromatin reorganization. To address this question, we utilized the same system described in Figure 5A to assess the immediate effects of acute ISWI depletion on CTCF binding. Of the 92,266 CTCF sites identified following MyoD nuclear localization, 85% (78,147) were constitutive, whereas 15% (14,119) showed MyoD-dependent recruitment (Figure 7D-E). Notably, only 14% (1,964) of the MyoD-dependent CTCF colocalized with MyoD, suggesting that MyoD likely facilitates CTCF recruitment indirectly. Interestingly, acute depletion of ISWI did not affect constitutive CTCF binding, but significantly disrupted MyoD-dependent CTCF binding (Figure 7F). Together, our data from the acute depletion system indicate that ISWI is dispensable for initial genomic binding of MyoD and preexisting CTCF, but is required for MyoD-dependent recruitment of cBAF, CBP, and CTCF.

## Discussion

In this study, we demonstrate that ISWI ATPases Snf2l and Snf2h are partially redundant and indispensable for embryonic development of muscle and BAT. ISWI is essential for myogenesis and adipogenesis of progenitor cells, as well as for LDTF-mediated cell fate transition. Acute depletion of ISWI reveals a distinct molecular requirement for ISWI in chromatin reorganization during cell fate transition. While ISWI is largely dispensable for initial binding of LDTFs, as exemplified with MyoD, it is critical for subsequent *de novo* recruitment of cBAF chromatin remodeler and architectural factor CTCF. Our findings suggest that global reduction of MyoD and CTCF binding in ISWI stable KO cells is likely an indirect consequence of altered chromatin organization. Together, our findings identify ISWI as a critical chromatin organizer that bridges LDTF binding to the *de novo* assembly of functional regulatory hubs during cell fate transition.

### Snf2h/Snf2l redundancy in differentiation & development

Several studies have reported important roles of Snf2h and Snf2l during mouse development. While constitutive *Snf2h* KO mouse embryos die at the peri-implantation stage, *Snf2l* KO mice can survive with relatively mild phenotypes, including increased brain size ^21,22^. During cerebellar development, Snf2h is required for the growth of Purkinje and granule cell progenitors, a function that can be partially compensated by the increased levels of Snf2l in the absence of Snf2h ^31^. In contrast, during lens development, differentiation defects in Snf2h-deficient precursor cells were not compensated by Snf2l ^23^. Consistent with these biological context-specific redundancy, our conditional KO models, *Snf2h*^f/f^;Myf5-Cre and *Snf2l*^-/-^;*Snf2h*^f/f^;*Myf5-Cre*, reveal distinct requirements of Snf2h and Snf2l during development of muscle and BAT. In BAT, *Snf2l/Snf2h* dKO embryos exhibited more severely reduced tissue mass compared to *Snf2h* single KO embryos. In line with these results, we observed increased *Snf2l* expression in *Snf2h* single KO BAT. In contrast, *Snf2h* single KO embryos already showed severe defects in muscle development. Notably, deletion of *Snf2h* alone is sufficient to block myogenesis and adipogenesis of isolated progenitor cells in culture, without noticeable upregulation of *Snf2l* expression. Together, these findings suggest lineage-selective functional redundancy between Snf2h and Snf2l during mouse embryonic development.

### Distinguishing primary and secondary effects of ISWI loss

By comparing the effects of stable KO versus acute depletion of ISWI, our study provides previously unrecognized mechanistic insights into the role of ISWI during cell fate transition. In ISWI stable KO cells, we observed a global collapse of the chromatin landscape, evidenced by widespread disruption of MyoD, cBAF, and CTCF binding, which leads to a failure of LDTF-driven myogenesis and adipogenesis. On the other hand, acute depletion of ISWI enabled us to distinguish primary and secondary effects caused by ISWI loss. MyoD genomic binding remained largely intact, yet the recruitment of chromatin remodeler cBAF was severely compromised. Prolonged depletion of ISWI further highlights the distinct effects. While MyoD binding progressively reduced over 48h, loss of cBAF binding was immediate and reached the maximum within 3h of ISWI depletion (Figure 5G). These results suggest that widespread defects in ISWI stable KO cells are likely secondary consequences of accumulated chromatin disorganization.

Our data also provides a refined assessment of the previously reported relationship between ISWI and CTCF. Earlier studies, using stable KO or knockdown approaches in relatively static cellular contexts such as mouse embryonic stem cells and human cancer cells, have identified ISWI as an essential regulator of CTCF genomic binding and nucleosome organization ^26,28^. In these systems, loss of ISWI caused a global reduction in CTCF binding. A more recent study using time-course SNF2H degradation in human blood cancer cell lines showed a gradual decrease in CTCF binding over several hours, with maximum effects observed between 6h and 24h after depletion ^32^. Using 3h acute depletion in the context of LDTF-initiated cell fate transition, we found that ISWI is required for establishing LDTF-dependent CTCF binding, while its absence does not exert immediate effect on constitutive CTCF binding. Together, these findings lead us to propose a model in which ISWI primarily supports LDTF-induced chromatin reorganization by facilitating recruitment of downstream regulators, including the chromatin remodeler cBAF and the architectural factor CTCF (Figure 7G).

### Distinct roles of ISWI and cBAF

A growing number of studies have reported a crucial role of cBAF in maintaining TF binding and chromatin accessibility in various cellular contexts. Acute catalytic inhibition (hereafter referred to as Brg1i) or degradation of the cBAF ATPases Brg1 and Brm leads to immediate loss of TF binding on chromatin. In mouse embryonic stem cells, Brg1i displaces the pluripotency TF OCT4 within 30 min ^33^. Similarly, in human neuroblastoma cells, Brg1i uniformly disrupts binding of adrenergic core regulatory TFs, HAND2, MYCN, ASCL1, ISL1, PHOX2B, and EBF3, within 1h ^34^. Treating prostate cancer cells with a PROTAC degrader of Brg1 and Brm, AU-15330, causes rapid loss of AR and FOXA1 binding within 1h ^35^. The cBAF-dependent genomic targeting also extends to *de novo* TF binding events. In breast cancer cells, Brg1i reduces GATA3 and FRA1 binding on GATA3-pioneered genomic regions ^36^. Our recent work demonstrates that Brg1i disrupts *de novo* binding of MyoD and GR ^27^. Together, these studies establish cBAF as a chromatin remodeler that is continuously required for both maintenance and acquisition of TF binding.

In contrast, our findings reveal a distinct role of ISWI during cell fate transition. Acute depletion of ISWI does not immediately displace MyoD from chromatin, but instead disrupts subsequent recruitment of cBAF. Although cBAF is widely perceived as the primary chromatin remodeler directly recruited by LDTFs to promote chromatin opening, our data identify ISWI as a prerequisite for LDTF-dependent cBAF recruitment. Together with existing literature, our study suggests that LDTF-driven chromatin reorganization is coordinated through distinct organizational layers, with ISWI establishing structural basis by regulating nucleosome spacing and cBAF subsequently generating chromatin accessibility required for LDTF-driven cell fate transition.

## Materials and Methods

### Antibodies and plasmids

All antibodies and plasmids used in this study are described in Supplementary Tables 1 and 2.

### Mouse strains and housing

To generate ISWI conditional KO (*Snf2l*^-/-^*;Snf2h*^f/f^) mice, *Snf2h*^f/f^ mice were crossed with *Snf2l*^-/-^mice ^22,23^. *Snf2l*^-/-^*;Snf2h*^f/f^ mice were crossed with *Myf5-Cre* (Jackson no. 007893) or *R26-CreERT2* (Jackson no. 008463) to generate *Snf2l*^-/-^*;Snf2h*^f/f^;*Myf5-Cre* mice or *Snf2l*^-/-^*;Snf2h*^f/f^;*CreER* mice. Mice were housed in a room with a controlled temperature of 22 °C and humidity of 45-65% under a 12-hour light and 12-hour dark cycle. All mouse work was approved by the Animal Care and Use Committee of NIDDK, NIH.

### Immortalization of primary brown preadipocytes and adipogenesis assay

Primary brown preadipocytes were isolated from interscapular BAT of newborn pups and were immortalized by infecting cells with retroviruses pBABE-neo largeT cDNA expressing SV40T as described ^37^. The immortalized cells were routinely cultured in growth medium (DMEM plus 10% FBS). For adipogenesis assays, preadipocytes were plated at a density of 1 × 10^5^ per well of 6-well plates in growth medium 4 days before induction of adipogenesis. At day 0, cells were fully confluent and were treated with differentiation medium (DMEM plus 10% FBS, 0.1 μM insulin, and 1 nM T3) supplemented with MDI (0.5 mM IBMX, 1 μM dexamethasone, and 0.125 mM indomethacin). Two days later, cells were changed to the differentiation medium. Cells were replenished with fresh medium at 2-day intervals. Fully differentiated cells were either stained with Oil Red O or subjected to gene expression analyses by qRT-PCR or RNA-Seq.

For adipogenesis assays in cells expressing WT and enzyme-dead SNF2H (Figure 2), immortalized *Snf2l*^-/-^;*Snf2h*^f/f^;*CreER* preadipocytes were infected with retroviruses expressing either pMSCVhygro-SNF2H(WT)-AID or pMSCVhygro-SNF2H(K211R)-AID. Following infection, cells were selected with 200 µg/ml hygromycin B (Sigma, #H3274). To delete endogenous *Snf2h*, cells were treated with 1 mM (Z)-4-Hydroxytamoxifen (4OHT), and then subjected to adipogenic differentiation.

For LDTF-stimulated adipogenesis assays (Figure 3), cells were infected with lentiviral Dox-inducible expression constructs (pCW57.1puro) encoding 3xT7-C/EBPα or 3xT7-PPARγ. Following infection, cells were selected with 2 µg/mL puromycin. Fully confluent WT and ISWI KO cells were treated with 1 mg/ml Dox to induce C/EBPα or PPARγ expression, together with MDI or 1mM rosiglitazone (Rosi) to promote adipogenic differentiation.

### Isolation of muscle satellite cells and myogenesis assay

Isolation of muscle satellite cells was carried out as described ^38^. Briefly, skeletal muscle was dissected from the hindlimbs of 8-week-old mice and minced with dissection scissors to small pieces (approx. 1 mm^3^). Minced muscle was enzymatically digested in Muscle Dissociation Buffer (1.5 mg/ml collagenase type 2 (Sigma, #C6885), 10 mM CaCl_2_ in HamF10 medium (ThermoFisher), no serum) for 1h with gentle swirling at 15 min intervals. Digested muscle tissue was suspended in two volumes of Wash Medium (HamF10 plus 10% horse serum), passed through a 40 μm strainer, and the mononuclear cells were recovered as a pellet by centrifugation (500 x g, 5 min). To collect muscle satellite cells, the mix of skeletal muscle cells was pre-plated on uncoated culture dishes for 3 hours, and then non-adherent cells were collected and plated on collagen-coated dishes (Corning). The muscle satellite cells were further expanded in the satellite cell growth medium (HamF10 plus 20% FBS, 5 ng/ml fibroblast growth factor-4 (FGF-4, Sigma, #F8424)). Myogenesis was induced when cells reached 80-90% confluency by replacing growth medium with differentiation medium (HamF10 plus 2% horse serum) and maintaining cells in differentiation medium for three days.

For the LDTF-stimulated myogenesis assay (Figure 3), immortalized *Snf2l*^-/-^;*Snf2h*^f/f^;*CreER* preadipocytes were infected with lentivirus expressing pCW57.1puro-MyoD-3xT7. Following infection, cells were selected with 2 µg/mL puromycin. To delete endogenous *Snf2h*, cells were treated with 1 µM 4OHT. WT and ISWI KO cells were cultured in growth media for 2 days before induction of myogenesis. Differentiation was initiated by switching to a medium consisting of DMEM supplemented with 2% horse serum (#H1270, Millipore-Sigma) and 1 μg/ml Dox. The medium was replenished every two days.

### *Snf2h* knockdown and KO by CRISPR-Cas9

For shRNA-mediated knockdown, individual shRNAs targeting mouse *Snf2h* (TRCN0000084430, sh*Snf2h*-#2; TRCN0000084432, sh*Snf2h*-#4) were obtained from Sigma Mission shRNA library (Sigma-Aldrich). Immortalized preadipocytes or C2C12 myoblasts were infected with lentiviral shRNAs targeting *Snf2h* or control virus alone for 24h. Infected cells were selected with puromycin (2 μg/mL) for 4 days before performing further experiments.

*Snf2h* single KO C2C12 cells were generated using CRISPR-Cas9. C2C12 cells were infected with LentiCRISPRv2-puro vector (Addgene #52961) expressing Cas9 and a sgRNA targeting exon 6 of *Snf2h* described previously ^26^. The infected cells were selected with puromycin (2 μg/ml) and cultured in growth media before performing further experiments.

### Acute depletion of SNF2H and nuclear translocation of MyoD in preadipocytes

To generate a cell line permitting acute ISWI depletion, *Snf2l*^-/-^;*Snf2h*^f/f^;*CreER* preadipocytes were infected with lentivirus expressing pLV-EF1a-IRES-puro-SNF2H-dTAG-2xHA. Following infection, cells were selected with 2 µg/ml puromycin and then treated with 1 µM 4OHT to delete endogenous *Snf2h*, generating *Snf2l*^-/-^;*Snf2h*^-/-^(ISWIKO);SNF2H-dTAG preadipocyte. For acute degradation of SNF2H-dTAG, dTAG-13 (Tocris Bioscience, #6605) was used at a final concentration of 0.5 µM.

ISWIKO;SNF2H-dTAG cells were further infected with retroviruses expressing pWZLhygro-MyoD-ER(T)-3xT7 ^27^. Infected cells were selected with 200 µg/ml hygromycin B (Sigma, #H3274) and maintained in growth medium. Nuclear translocation of MyoD-ER was induced by treating cells with 400 nM 4OHT.

### Western blot

Nuclear proteins were extracted from preadipocytes. Cells were washed with cold PBS, resuspended in buffer A (10 mM HEPES, pH 7.9, 1.5 mM MgCl_2_, 10 mM KCl, and 0.1% NP40) supplemented with protease inhibitors (Roche), 0.5 mM DTT and 0.2 mM phenylmethylsulfonyl fluoride (PMSF), and incubated on ice for 10 min. After centrifugation at 1,000g, nuclear proteins were extracted in buffer C (20 mM HEPES, pH 7.9, 1.5 mM MgCl_2_, 420 mM NaCl, 0.2 mM EDTA, and 25% glycerol) supplemented with protease inhibitors, 0.5 mM DTT and 0.2 mM PMSF. Nuclear extracts were separated using 4–15% Tris-Glycine gradient gels (Bio-Rad Laboratories), and transferred to a PVDF membrane using the iBlot 2 Gel Transfer Device (Life Technologies). The membranes were probed using specific antibodies.

### RNA isolation and RNA-Seq library preparation

Total RNA was extracted using TRIzol (Life Technologies) and subjected to mRNA purification using the NEBNext Poly(A) mRNA Magnetic Isolation Module (NEB, E7490), following the manufacturer’s instructions. Purified mRNAs were reverse-transcribed into double-stranded cDNA and subjected to sequencing library construction using the NEBNext Ultra™ II RNA Library Prep Kit for Illumina (NEB, E7770) with unique dual index primer pairs (NEB, E6440) according to the manufacturer’s instructions. RNA libraries were sequenced on Illumina HiSeq 3000.

### Chromatin immunoprecipitation (ChIP) and ChIP-Seq library preparation

Chromatin immunoprecipitation (ChIP) and ChIP-Seq were done as described ^3^. In summary, cells were cross-linked with 1% formaldehyde for 10 minutes and then quenched using 125 mM glycine. Nuclei were isolated from ten million cells using hypotonic buffer containing 5 mM PIPES (pH 7.5), 85 mM KCl, 1% NP-40, and protease inhibitors, and resuspended in 1 mL of a buffer composed of 50 mM Tris-HCl (pH 8.0), 10 mM EDTA, 0.1% SDS, and protease inhibitors, followed by sonication. The sheared chromatin was clarified by centrifugation at 13,000 g for 10 minutes at 4°C. The supernatant was transferred to a new tube and supplemented with 1% Triton-X100, 0.1% sodium deoxycholate, and protease inhibitors. Each ChIP reaction included 4 to 10 µg of target antibodies, 20 ng of spike-in chromatin (Active Motif, #53083), and 2 µg of spike-in antibody (anti-H2Av, Active Motif, #61686), incubated on a rotator at 4°C overnight. ChIP samples were then mixed with 50 μL of prewashed Protein A Dynabeads (ThermoFisher) and incubated for 3 hours at 4°C. The beads were collected using a magnetic rack and washed twice with RIPA buffer, RIPA buffer containing 300 mM NaCl, cold LiCl buffer, and PBS. ChIP and input samples were added to 100 μL of ChIP elution buffer containing 0.1 M NaHCO3, 1% SDS, and 20 μg of Proteinase K, and incubated at 65°C overnight. Purified DNA using the QIAquick PCR purification kit (Qiagen). For ChIP-Seq library preparation, the purified DNA was utilized to create libraries using the NEBNext® Ultra™ II DNA Library Prep kit alongside AMPure XP magnetic beads (Beckman Coulter). The quality and quantity of the libraries were evaluated with Tapestation and Qubit assays, and the libraries were paired-end sequenced on either the Illumina NovaSeq X or NextSeq 2000 systems.

### Computational analysis

#### RNA-Seq data analysis

Raw sequencing data were aligned to the mouse genome mm9 using STAR software ^39^. Reads mapped to exonic regions were used to calculate reads per kilobase per million (RPKM) as a metric for gene expression levels. Genes with exonic reads of RPKM > 1 were considered expressed. To compare gene expression levels in satellite cells (D0) and myoblasts (D3) (Fig. 2), before (D-3) and during (D2) adipogenesis (Fig. 2-S1), or between control and ISWI-deficient cells, a fold change cutoff of > 2 was applied to identify differentially expressed genes. Gene ontology (GO) analysis was done using DAVID, with the entire mouse genome as the background (https://david.ncifcrf.gov).

#### ChIP-Seq peak calling and motif analysis

Raw sequencing data were aligned to the mouse genome mm10, as well as the drosophila genome dm6 using Bowtie2 (v2.3.2) ^40^. To identify enriched regions from ChIP-Seq, we used SICER (v2) ^41^ with the following parameters: a window size of 50 bp, a gap size of 50 bp, and an FDR threshold of 10^-10^. To identify enriched TF motifs, we used the HOMER software (http://homer.ucsd.edu/homer/) ^42^.

#### Read count quantification and spike-in normalization

Read counts were quantified over a fixed set of genomic regions. Reads overlapping each region were calculated from the SICER output (islandfiltered.bed). Counts were first normalized by dividing by the total number of reads in each sample and multiplying by 1,000,000. These normalized values were then multiplied by their respective spike-in scaling factors derived from *Drosophila* dm6 reads. The scaling factor for each sample was calculated by dividing the dm6 read count of the sample with the lowest dm6 reads by that of each corresponding sample.

#### Heat maps

For heat map visualization, genomic regions were grouped into two categories based on fold changes in spike-in-normalized read counts in ISWI KO or depleted cells relative to control cells. Regions with >2-fold decrease were assigned to a “decreased” group, and remaining regions were classified as “intact”.

Heat maps were generated using deepTools (v3.5.1) ^43^. First, input-filtered alignment files were converted to normalized bigWig signal tracks using bamCoverage, with scaling factors obtained from *Drosophila* spike-in read counts applied to adjust differences in signal intensity across conditions. Signal enrichment was quantified using computeMatrix in reference-point mode, centered on peak regions, with signal calculated in ±5 kb windows around the peak center. The resulting matrices were visualized using plotHeatmap.

### Data Availability

The data that support this study are available from the corresponding author upon reasonable request. All ChIP-Seq and RNA-Seq datasets described in this study have been deposited in NCBI Gene Expression Omnibus under access #GSE318620 [https://www.ncbi.nlm.nih.gov/geo/query/acc.cgi?acc=GSE318620].

## Acknowledgements

We thank the National Heart, Lung, and Blood Institute DNA Sequencing and Genomics Core and the National Institute of Arthritis and Musculoskeletal and Skin Diseases Genomic Technology Section for next-generation sequencing. This work was supported by the Intramural Research Program of the National Institute of Diabetes and Digestive and Kidney Diseases (NIDDK), National Institutes of Health to K.G. A.I.S. was supported by NIH grant R01 GM147165.

## Author Contributions

Conceptualization, Y.-K.P., J.-E.L., and K.G.; Methodology, Y.-K.P., J.-E.L., A.I.S., and D.J.P.; Investigation, Y.-K.P. and K.G; Software, Formal Analysis, and Data Curation, J.-E.L. and W.P.; Writing – Original Draft, Y.-K.P. and J.-E.L.; Writing – Review & Editing, Y.-K.P., J.-E.L., and K.G.; Project Administration and Funding Acquisition, K.G.

## NIH Acknowledgement Disclaimer Statement

This research was supported by the Intramural Research Program of the National Institute of Diabetes and Digestive and Kidney Diseases (NIDDK) within the National Institutes of Health (NIH). The contributions of the NIH authors are considered Works of the United States Government. The findings and conclusions presented in this paper are those of the authors and do not necessarily reflect the views of the NIH or the U.S. Department of Health and Human Services.

## Figure legends

**Figure 2-S1.**
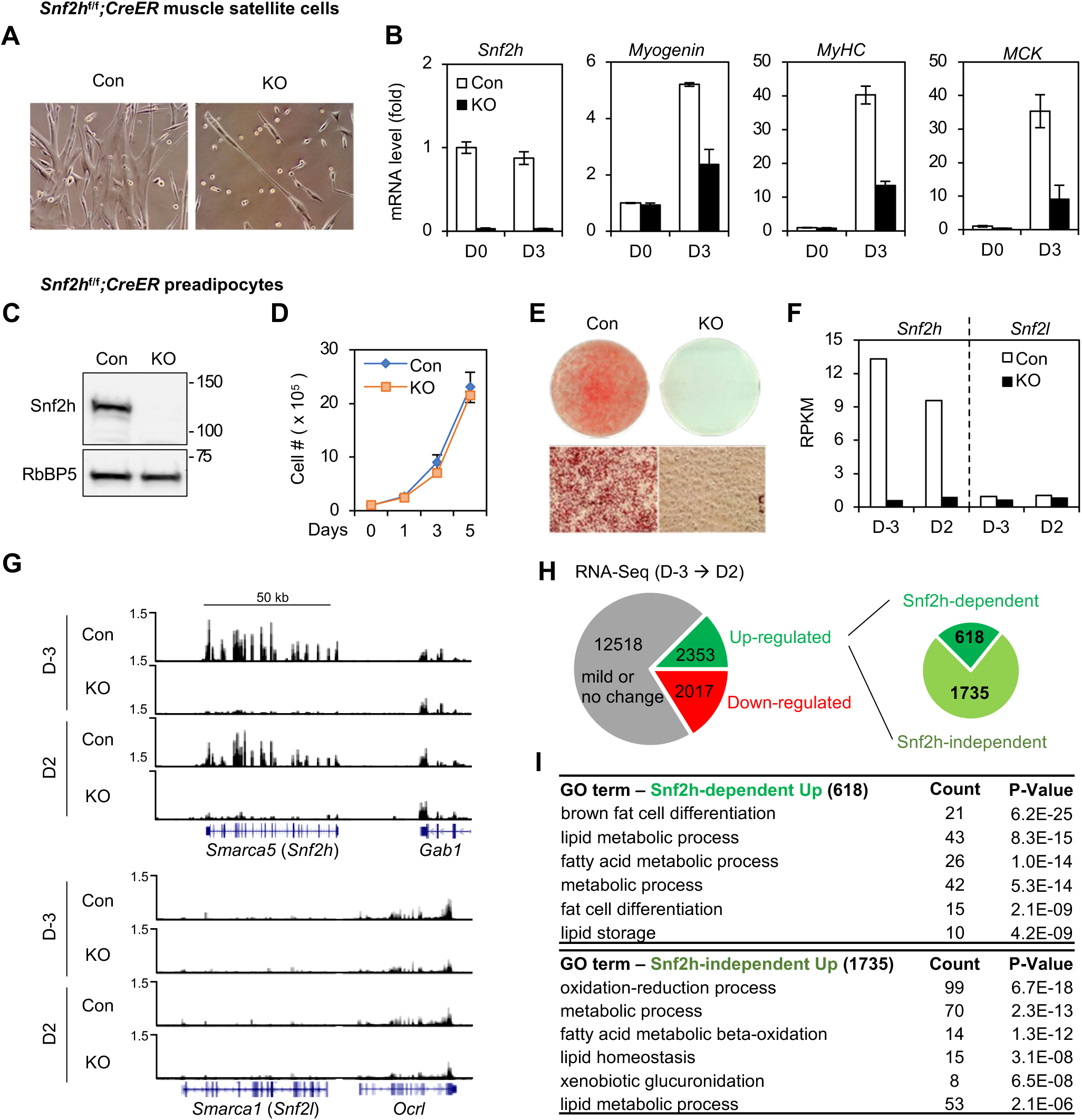
Snf2h is required for myogenesis and adipogenesis in culture. (**A-B**) Snf2h is required for myogenesis in culture. Muscle satellite cells isolated from *Snf2h*^f/f^;*CreER* mice were treated with 4OHT, followed by myogenesis assays. (**A**) Representative microscopic pictures of satellite cells at D3 of myogenesis. (**B**) qRT-PCR analysis of myocyte marker genes *Myog*, *Myh1*, and *Ckm* during myogenesis. (**C-E**) Snf2h is required for adipogenesis in culture. SV40T-immortalized *Snf2h*^f/f^;*CreER* preadipocytes were treated with 4OHT, followed by adipogenesis assays. (**C**) Western blot analysis of Snf2h in preadipocytes. RbBP5 was used as a loading control. (**D**) Deletion of *Snf2h* does not affect cell growth rates. 1 x 10^5^ preadipocytes were plated and cumulative cell numbers were determined for 5 days. (**E**) Oil Red O staining at D7 of adipogenesis. (**F-I**) RNA-Seq analysis before (D-3) and during (D2) adipogenesis. (**F**) Expression of *Snf2h* and *Snf2l* at D-3 and D2. RPKM values indicate gene expression levels. (**G**) Genome browser view of *Snf2h* (upper panel) and *Snf2l* (lower panel) loci at D-3 and D2. (**H**) Pie chart depicts Snf2h-dependent and - independent up-regulated genes as well as down-regulated genes from D-3 to D2 of adipogenesis. The cut-off for differential expression is 2-fold. (**I**) Gene ontology (GO) analysis of 618 Snf2h-dependent and 1735 Snf2h-independent up-regulated genes defined in (**H**).

**Figure 2-S2.**
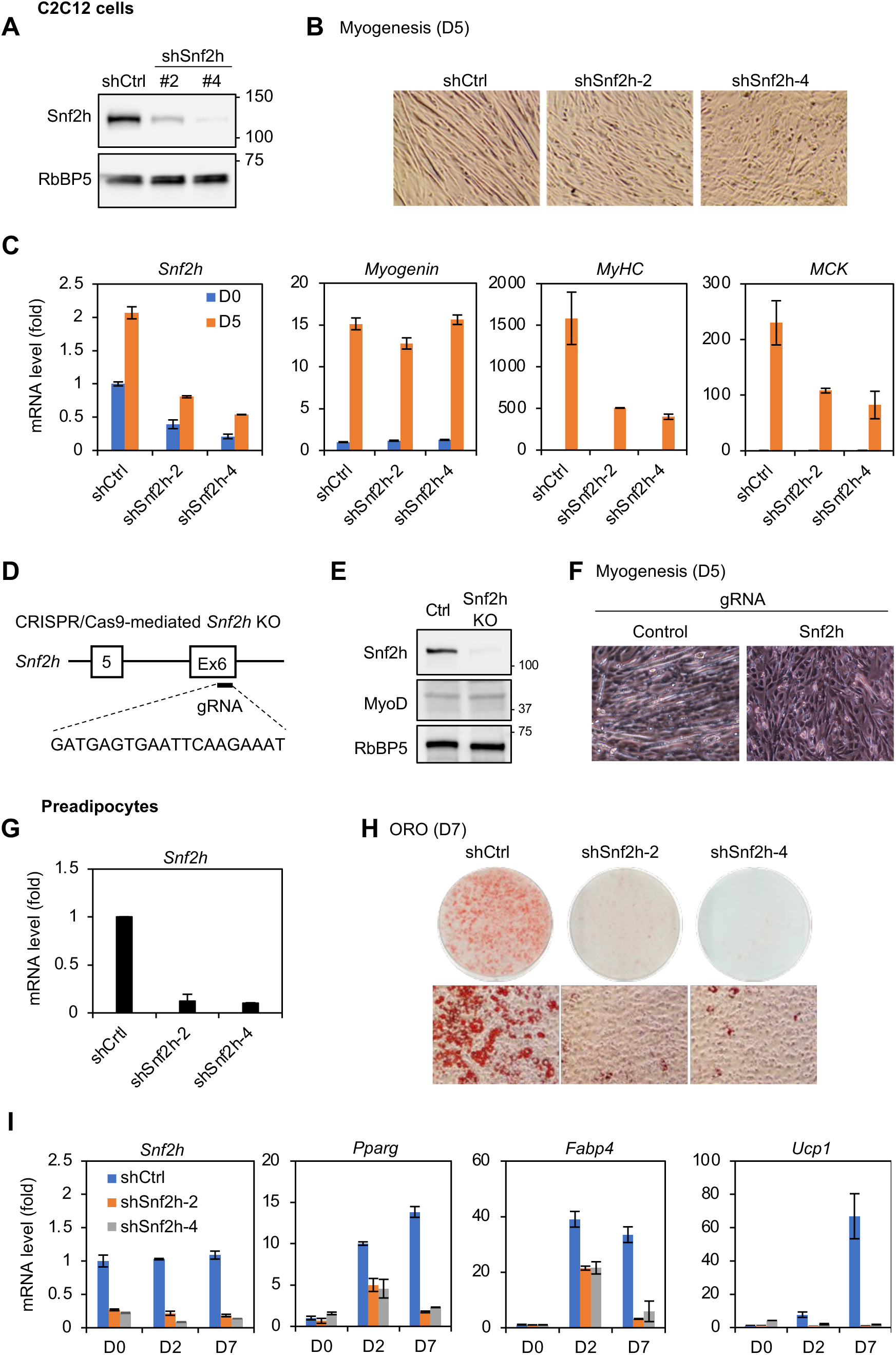
Snf2h depletion prevents myogenesis and adipogenesis in culture. (**A-C**) C2C12 cells were infected with lentiviral vector expressing control (Ctrl) or *Snf2h* knockdown shRNA (sh*Snf2h*), followed by myogenesis assays. (**A**) Western blot analysis of Snf2h in C2C12 cells. (**B**) Representative microscopic pictures at D5 of myogenesis. (**C**) qRT-PCR analysis of myocyte marker genes *Myog*, *Myh1*, and *Ckm* during myogenesis. (**D-F**) Myogenesis in CRISPR/Cas9-mediated *Snf2h* KO C2C12 cells. (**D**) Schematic of generating *Snf2h* KO C2C12 cells using a gRNA. (**E**) Western blot analysis of Snf2h and MyoD protein levels in control and *Snf2h* KO C2C12 cells. RbBP5 was used as the loading control. (**F**) Microscopic images at D5 of myogenesis. (**G-I**) Preadipocytes were infected with lentiviral vector expressing control or *Snf2h* knockdown shRNA, followed by adipogenesis assays. (**G**) mRNA levels of *Snf2h* in preadipocytes. (**H**) Oil Red O staining of differentiated cells at D7. (**I**) qRT-PCR analysis of adipocyte marker genes *Pparg*, *Fabp4*, and *Ucp1* gene expression at indicated days.

**Figure 3-S1.**
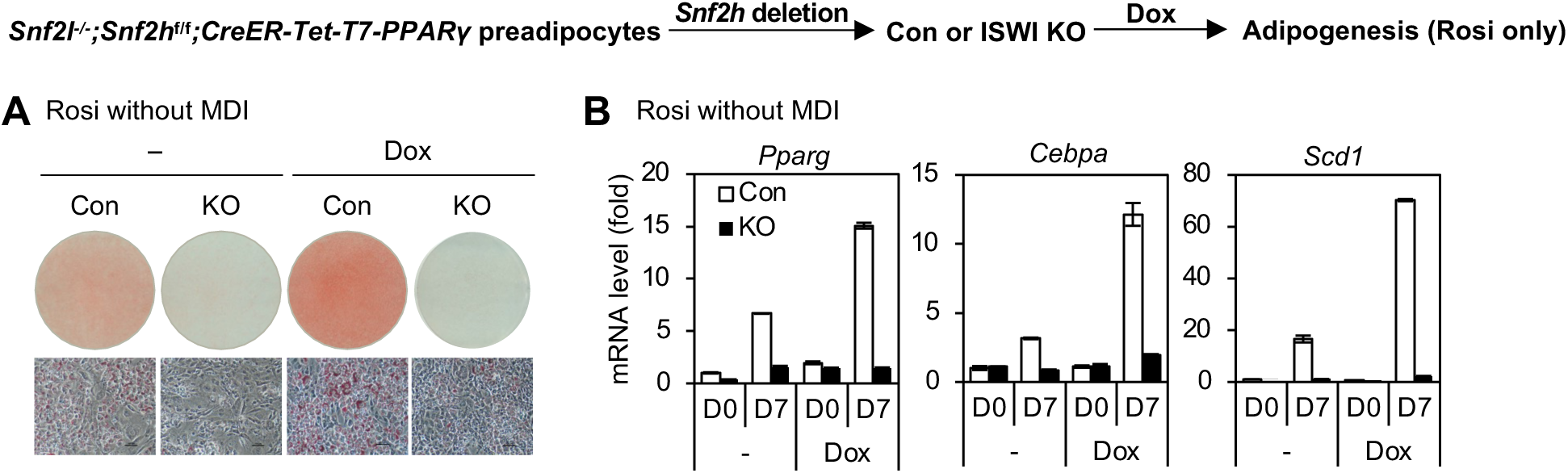
Stable KO of ISWI prevents PPARγ ligand-stimulated adipogenesis. *Snf2l*^-/-^;*Snf2h*^f/f^;*CreER* preadipocytes were infected with lentiviruses expressing Dox-inducible T7-PPARγ. Cells were treated with 4OHT to delete *Snf2h*, followed by adipogenesis assay with 1 μM Rosi only in the presence of Dox for 7 days. (**A**) Oil Red O staining at D7 of adipogenesis. (**B**) qRT-PCR analysis of adipocyte marker genes *Pparg, Cebpa,* and *Scd1* at D0 and D7 of adipogenesis.

**Figure 4-S1.**
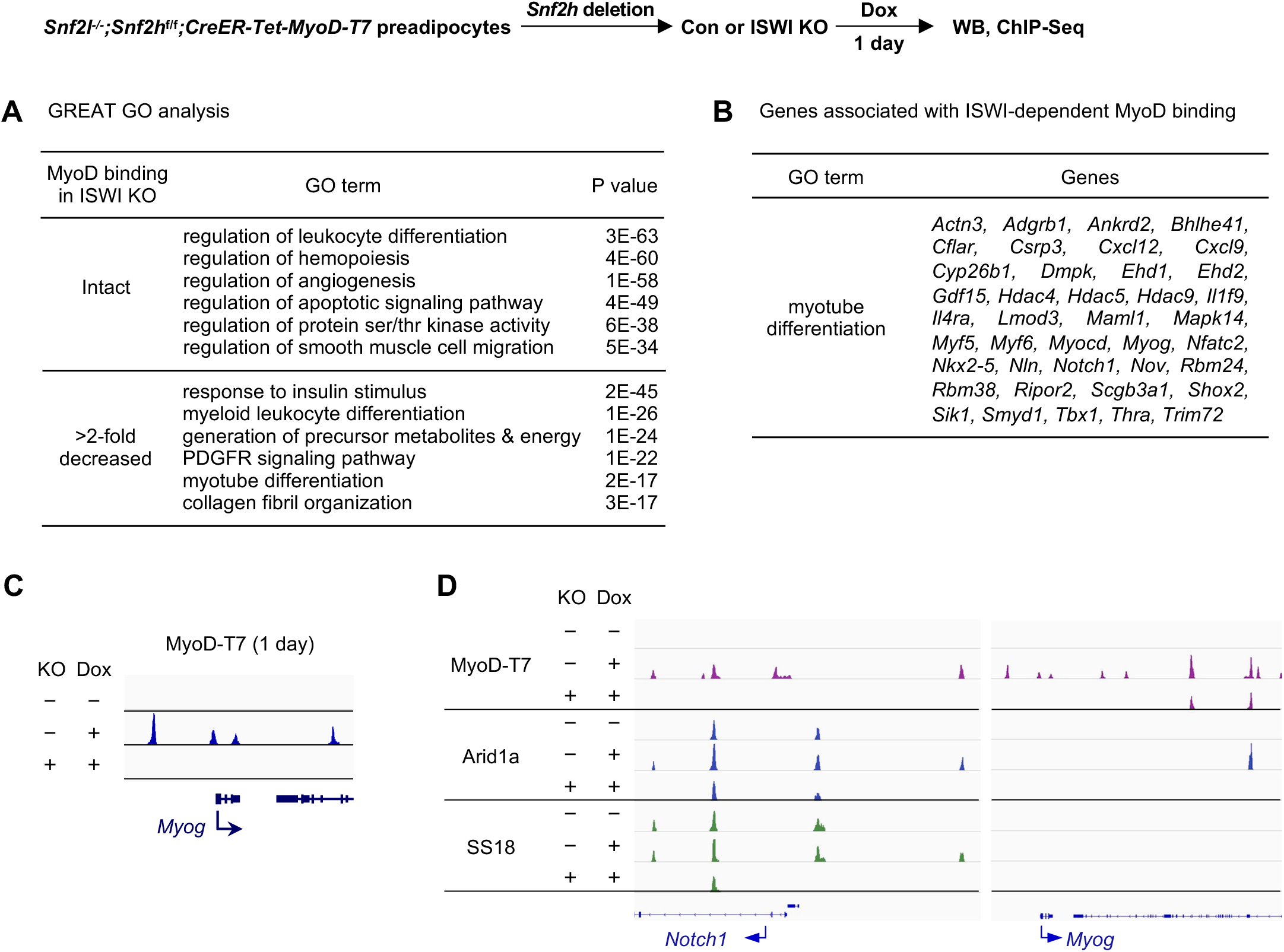
Stable KO of ISWI disrupts *de novo* binding of MyoD and cBAF on chromatin. *Snf2l*^-/-^;*Snf2h*^f/f^;*CreER* preadipocytes were infected with lentiviruses expressing Dox-inducible MyoD-T7 (MyoD). Cells were treated with 4OHT to delete *Snf2h*, followed by 1 day Dox treatment to induce MyoD expression. Sub-confluent cells were collected for ChIP-Seq analyses. (**A**) GREAT GO analysis of intact MyoD (18,727) and reduced MyoD (17,941) sites defined in Figure 4B. (**B**) List of genes associated with ISWI-dependent MyoD binding and involved in myotube differentiation, as identified in (**A**). (**C**) Genome browser view of MyoD binding on *Myog* and neighboring gene loci. (**D**) Genome browser view of MyoD, Arid1a and SS18 binding around *Notch1* and *Myog* loci.

**Figure 5-S1.**
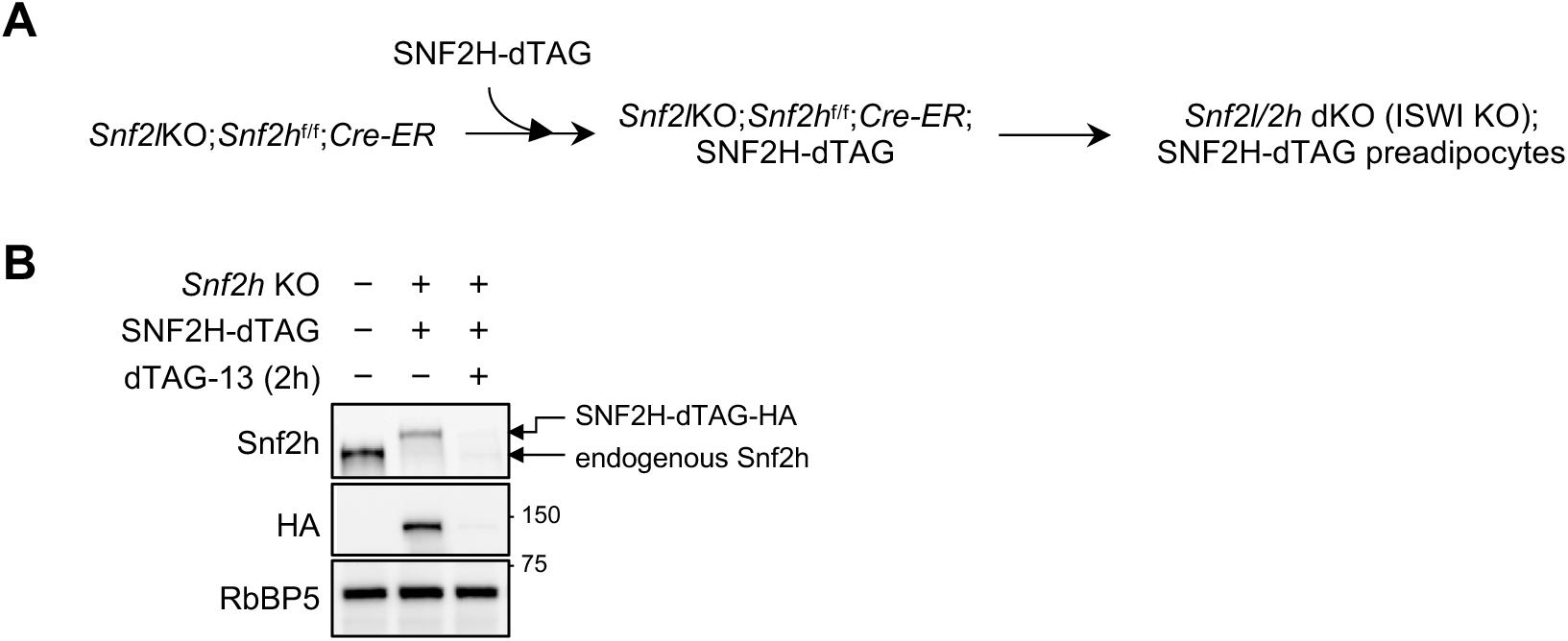
Generation of cell lines for acute SNF2H depletion using dTAG system. (**A**) Schematic illustrating the generation of ISWI KO; SNF2H-dTAG preadipocytes. (**B**) Western blot analysis of endogenous Snf2h and ectopic SNF2H-dTAG-HA expression after dTAG-13 treatment for 2h. RbBP5 was used as a loading control.

**Figure 6-S1.**
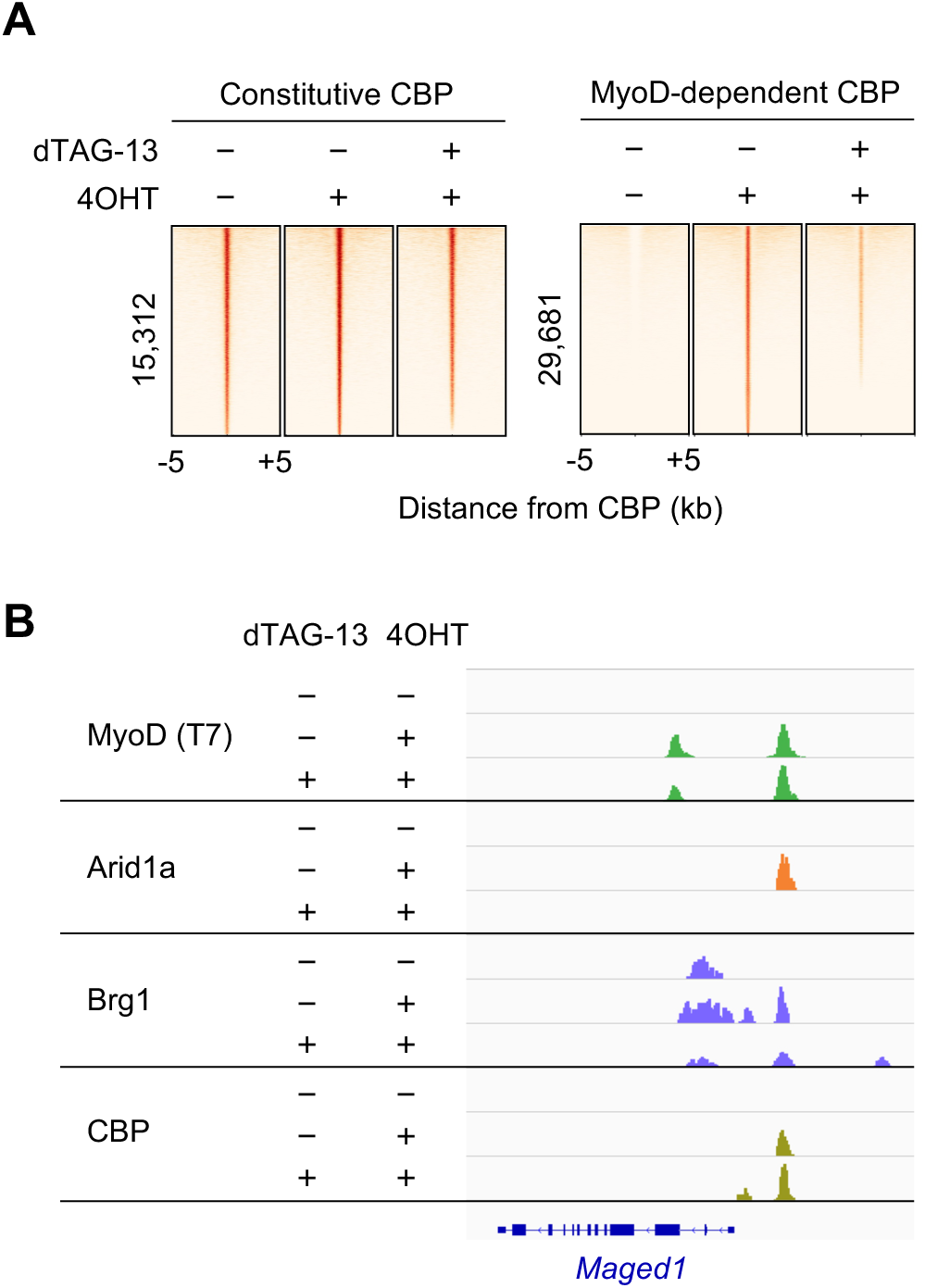
Acute depletion of ISWI disrupts *de novo* cBAF binding. (**A**) Acute depletion of ISWI disrupts MyoD-dependent, but not constitutive, CBP binding. Heat maps illustrate changes in CBP binding intensity on constitutive or MyoD-dependent CBP sites. (**B**) Genome browser view of MyoD, Arid1a, Brg1 and CBP binding around the *Maged1* locus.

